# Soil detrital inputs increase stimulate bacterial saprotrophs with different timing and intensity compared to fungal saprotrophs

**DOI:** 10.1101/266684

**Authors:** Nicole Sukdeo, Ewing Teen, P. Michael Rutherford, Hugues B. Massicotte, Keith N. Egger

## Abstract

Soils contain microbial inhabitants that differ in sensitivity to anthropogenic modification. Soil reclamation relies on monitoring these communities to evaluate ecosystem functions recovery post-disturbance. DNA metabarcoding and soil enzyme assays provide information about microbial functional guilds and organic matter decomposition activities respectively. However bacterial communities, fungal communities, and enzyme activities may not be equally informative for monitoring reclaimed soils. We compared effects of disturbance regimes applied to forest soils on fungal community composition, bacterial community composition, and potential hydrolase activities (N-acetyl-β-D-glucosaminidase, acid phosphatase, and cellobiohydrolase) at two times (14 days and 5 months post-disturbance) and depths (LFH versus mineral soil). Using disturbance versus control comparisons allowed us to identify genus-level disturbance-indicators and shifts in hydrolase activity levels. We observed declines in disturbed LFH fungal biomass (ergosterol) and declines in ectomycorrhizal fungi abundance across all disturbed samples, which prompted us to consider necromass-induced (fungal, root) saprotroph increases as disturbance indicators. Fungal community composition strongly shifted away from ecotmycorrhizal dominance to saprotroph dominance (i.e. increased *Mortierella*, and *Umbelopsis*) in disturbed plots at 5 months, while bacterial community composition did not shift to distinguish control plots from disturbed ones at either sampling time. Soil potential hydrolase data mainly indicated that mixing LFH material into mineral soil increases the measured activity levels compared to control and replaced mineral soil. Bacterial saprotrophs previously associated with mycelial necromass were detected across multiple regimes as disturbance indicators at 14 days post-disturbance. Our results confirm that ectomycorrhizal fungal genera are sensitive and persistently impacted by soil physical disturbances. Increases in saprotrophic bacterial genera are detectable 14 days pot-disturbance but only a few persist as disturbance indicators after several months. Potential hydrolase activities appear to be most useful for detecting the transfer of decomposition hotspots into mineral soils.

## Introduction

Physical disturbances of soil are an inevitable consequence of anthropogenic modifications to terrestrial ecosystems. Anthropogenic impacts on forest soils are a focus of many DNA metabarcoding studies that aim to characterize microbial responses to disturbance. Microbial communities in forest soils change because of various disruptions including fires (Glassman et al., 2015), logging (Hartmann et al., 2014; Kyaschenko et al., 2017; Leung et al., 2016), tree mortality from insect infestations (Saravesi et al., 2015; Stursová et al., 2014; Treu et al., 2014), and compaction (Hartmann et al., 2014), but timing and effect size of disturbance responses are different for fungi and bacteria (Hartmann et al., 2014, 2012; Stursová et al., 2014; Sun et al., 2017). Differences in how bacteria and fungi are distributed in soil pore space and within rhizospheres account for some differences in disturbance responses (Uroz et al., 2016). Boreal forest soils with dense overstory vegetation will have extensive biomass from ectomycorrhizal (EcM) fungal hyphae that physically associate with plant roots and explore soil pore space for nutrient acquisition (Churchland and Grayston, 2014). Disruptions to EcM networks by logging, tree mortality or other disturbances to carbon flow from roots stimulate rhizosphere saprotrophs that access nutrient pools consisting of root and fungal necromass (Fernandez et al., 2016; Lindahl et al., 2010). The Gadgil effect, which describes community dominance and increased organic matter decomposition by saprotrophic fungi when mycorrhizal abundance decreases, is a well-known disturbance response in forest systems (Fernandez and Kennedy, 2016; Gadgil and Gadgil, 1971), and has been demonstrated through amplicon-sequencing surveys of fungal communities in disturbed forest systems (Kyaschenko et al., 2017; Saravesi et al., 2015; Stursová et al., 2014).

Interpreting bacterial community shifts in disturbance contexts is challenging because of the paucity of trait-based frameworks for describing bacterial ecosystem services and response to environmental change (Barberán et al., 2014; Martiny et al., 2015). Bacteria occupy regions in rhizosphere and mycorrhizosphere soil compartments where they provide ecosystem services to plants and mycorrhizal fungi, or contribute to organic matter turnover as these sites generate detritus. Like soil fungi, bacteria colonize decomposing litter (Brabcová et al., 2016) and filamentous Actinobacteria can perform hyphal exploration (Jones et al., 2017). Bacteria can also inhabit biofilms (Baldrian, 2017), move through soil via attachment to exploratory fungal hyphae (Bravo et al., 2013; Simon et al., 2015), or through inherent motility repertoires.

Kuzyakov and Blagodatskaya (Kuzyakov and Blagodatskaya, 2015) define soil locations like rhizospheres, mycorrhizospheres, detrituspheres (i.e. decomposing litter), and aggregated biofilm communities as “hotspots” of microbial activity because they are locations where biogeochemical processes occur at elevated rates relative to bulk soil. Precise sampling of soil hotspots permits localized measurement of processes (i.e. respiration, sites of enzyme activity, increased microbial biomass), but does not inform how hotspots are distributed in soil volumes that are relevant to anthropogenic disturbance effects.

Mixing of soil horizons and rhizosphere/mycorrhizosphere disruption caused by disturbance of forest systems changes locations of detritus and substrate pools at a local scale. Microbial community profiling by DNA metabarcoding of soils sampled weeks, months, or years after disturbance may reveal community members that respond to this disturbance. LFH and mineral soil mixing (Macdonald et al., 2015; Soon et al., 2000) as well as topsoil stockpiling for future soil profile re-assembly (Rokich et al., 2000), are components of soil disturbance that have implications for the abundance and persistence of hot spots associated with organic matter turnover, or re-establishment of fungal-bacterial community assemblies that optimize mycorrhizal function. Therefore, soil community analysis via DNA metabarcoding can help to link microbial disturbance responses to different hotspot types and distributions caused by different soil handling strategies. While disruptions of EcM fungi in disturbed forests are consistently identifiable through ITS2 sequencing surveys of bulk LFH and mineral soils, (Kyaschenko et al., 2017; Stursová et al., 2014; Sukdeo et al., 2018; Sun et al., 2015) disturbance responses of bacteria are more variable (Bastida et al., 2017; Hartmann et al., 2012; Wilhelm et al., 2017).

The objective of this study was to examine the utility of fungal community profiles, bacterial community profiles, and soil potential hydrolase activities in monitoring forest soil disturbance. We used 16S and ITS2 Illumina-tag sequencing to compare bacterial and fungal communities between undisturbed sub-boreal forest soils and three soil disturbance regimes. The disturbance regimes emulate different soil reclamation strategies (i.e. approximate horizon replacement versus mixed LFH and mineral soil) but they all generate mycelial and root necromass inputs that are hotspots for microbial saprotrophs. We used Linear discriminant effect size analysis (LEfSe) (Segata et al., 2011) for disturbance regime versus control comparisons within sampling times to identify responsive fungal or bacterial genera, and determine whether these genera are enriched at 14 days and/or five months after soil manipulation. Soil potential hydrolase activities (specifically N-acetyl-β-D-glucosaminidase, acid phosphatase, and cellobiohydrolase) were also compared between treatments to infer whether enzymatic litter/detritus decomposition activities are detectable in bulk soil samples with recent introduction of detrital inputs from disturbance.

## Materials and Methods

### Site description, experimental design, and sample collection

We selected two field sites approximately 60 km northeast of Prince George, British Columbia (Site 1: 54.2007 N, -123.1600 W; Site 2: 54.1992 N, -123.1606 W), within “Moist Cool Sub-boreal Spruce biogeoclimatic subzones (SBSmk1) of the sub-boreal spruce (SBS) biogeoclimatic zone (Beaudry et al., 1999). The sites were approximately 170 m apart from each other. Soils at both sites were classified as Eluviated Dystric Brunisols within the Canadian System of Soil Classification (SCWG, 1998). Soil pit descriptions for these field sites are previously described in Sukdeo et al. (2018). At each site, three replicate areas or blocks were selected with a minimum distance of 10 m between them, so each block could accommodate four 1 m × 2 m rectangular plots. Within a block, each plot was randomly assigned as a single technical replicate of the three soil disturbance configurations, or a control, undisturbed configuration (4 treatments in total). Disturbance regimes were applied to plots on May 12-14, 2014. A total of 24 field plots (4 treatments × 3 blocks/replicates × 2 sites) were established and soil samples were collected at 14 days (May 26, 2014) and 5 months (November 6, 2014) post-disturbance. Upper soil samples were collected to a depth of 10 cm to evaluate treatment effects on LFH material. Lower soil samples consisted of underlying mineral soils collected a maximum depth of 20 cm in plots. Treatment B (described below) soil was predominantly mineral in composition, collected to a depth of 10 cm, and classified as a lower soil sample.

The soil disturbance regimes were as follows: A = undisturbed plot (Control); B = 20 cm depth excavation, organic LFH layer manually mixed into underlying mineral soil; C = 20 cm depth excavation, LFH scraped off and sequestered (pile), lower mineral soil was manually mixed and immediately covered with sequestered LFH material; D = 20 cm depth excavation, LFH scraped off and stored on gardening textile for 5 months (semi-protected under adjacent canopy), lower mineral soil was manually mixed and left uncovered until November 6, 2014 (i.e. 5 months post-disturbance), when the LFH stockpile was placed back on the plot. The relevance of each disturbance regime to soil reclamation practices, details of soil sampling, and soil storage is summarized in Sukdeo et al. 2018.

### Soil ergosterol content

Ergosterol was isolated by KOH/methanol extraction and quantified as a proxy for fungal biomass (Seitz et al., 1977) by high performance liquid chromatography (Chiocchio and Matković, 2011; Newell et al., 1988). Kruskal-Wallis tests for distribution of ergosterol values between disturbance regimes within sampling depths at the 5-month time-point were previously reported in Sukdeo et al. 2018. This data is summarized in Supplementary Table 1. These analyses were run in R version 3.1.2 (R Core Development Team, 2014) and pairwise comparisons were made between treatments for significant (p < 0.05) results using the Holm-corrected Dunn’s test in R package PMCMR (Pohlert, 2014).

### Soil potential hydrolase activity assays

All substrates were purchased from Sigma-Aldrich (St. Louis, MO, USA). The methylumbelliferone (MUB)-linked substrates 4-MUB-phosphate (CAS 3368-04-5), 4-MUB-N-acetyl-β-D-glucosaminide (CAS 37067-30-4), and 4-MUB-β-D-cellobioside (CAS 72626-61-0) were used to assay for potential acid phosphatase (PHOS), N-acetyl-β-D-glucosaminidase (NAG), and cellobiohydrolase (CBH) activities, respectively. Soil homogenates were prepared by adding 10mL of 50 mM NaCH_3_COO (acetate) buffer (pH 5.2) to 1g soil (wet weight) and vortexing for 30 min (3000 rpm, using a Fisher pulse vortex mixer, model number 945420) at ambient temperature. Four replicate measurements were made for every sample in the assay. 128 μl of soil suspension were added to 32 μl of 4-MUB-linked substrate (1.25 mM stock concentration, final concentration in assay was 250 μM) in flat bottom, 96-well black microplates (part number 7605; Thermo Scientific™, Waltham, MA, USA). Substrate and buffer autofluorescence were measured by adding 32 μl of substrate solution and 32 μl of acetate buffer, respectively, to 160 μl of acetate buffer. A standard curve was generated using 4-methylumbelliferone (4-MUB) solutions diluted in acetate buffer (final volume of dilutions in microplates was 160 μl) with the following 4-MUB concentrations ranging from 0 μM to 50 μM. Soil quench curve data was collected using 132 μl of homogenates from single upper and lower soil samples from Treatment A (control) and a 4-MUB concentration range identical to of the standard curve. Microplates were incubated at room temperature for 2 hours in the dark. Plates were read with a FilterMax F5 MultiMode Microplate Reader and the SoftMax^®^ Pro 6.3 software (Molecular Devices, Sunnyvale, CA, USA) using a 360 nm excitation filter and a 460 nm emission filter.

Potential hydrolase activities were calculated using the formulas for emission coefficients, quench coefficients, and activity (in units nmol h^−1^ g^−1^) as described in (German et al., 2011). Calculated hydrolase activities were converted so that units reflected activity in dry mass equivalents of soil samples, using gravimetric water content measurements determined by mass loss following oven-drying at 105°C for 24 hours (Kalra and Maynard, 1991).

### DNA extraction, fungal ITS2 and bacterial 16S V4 amplicon preparation

A single 0.25 g sub-sample for each treatment replicate at all sampling times was used for DNA extractions using the PowerSoil DNA Isolation Kit (Mo Bio Laboratories, Carlsbad, CA, USA) according to manufacturer’s instructions. Primer sequences and thermal cycling conditions for amplifying fungal ITS2 regions were for the ITS86F/ITS4 primer pair as described in (Vancov and Keen, 2009). Primer sequences and thermal cycling conditions for amplifying bacterial 16S V4 regions were for the 515F/806R primer pair as described in (Caporaso et al., 2012). Prior to PCR amplification, 10 ng of template DNA from each sample was pre-incubated with 0.5 μg BSA (New England Biolabs) for 10 minutes at 95 °C prior to the addition of master mix containing primers, 5 PRIME Hot Master mix (2.5 X stock) and nuclease free water (IDT). The final concentration of PCR components in a 25 μl reaction volume (including template and BSA) were 1X 5 PRIME Hot Master Mix and primers (bacterial or fungal) at 200 nM final concentration. For each sample triplicate, 25 μL PCR’s were pooled and the amplicons were extracted from 1% agarose, 0.5X TBE gels after electrophoresis using the GeneJET Gel Extraction and DNA Cleanup Micro Kit (ThermoFisher Scientific) and eluted in 20 μL of elution buffer.

DNA concentrations of the purified amplicons were determined with a Qubit Fluorometric assay (ThermoFisher Scientific) using the dsDNA BR assay kit. DNA concentrations were manually adjusted to 2 ng/ml and pooled and submitted for sequencing to the Génome Québec Innovation Centre’s Massively Parallel Sequencing Services unit (McGill University, Montreal, QC, Canada). The amplicon pools were sequenced, using 300 nt paired-end chemistry (ITS2) or 150 nt paired-end chemistry (16S) on an Illumina MiSeq instrument. Raw sequence data were deposited at the National Centre for Biotechnology Information (NCBI, US) Sequence Read Archive under the Study accession number SRP107088 and Bioproject accession number PRJNA38668. Supplementary Text 1 describes data processing steps and Supplementary Text 2 summarizes the number of high-quality sequences, OTU bins, and genus-classified bins for bacterial 16S amplicon sequence libraries. ITS2 amplicon sequence library data processing steps and summaries of high-quality sequences, OTU bins, and genus-classified bins are described in Sukdeo et al. 2018. Hierarchical cluster (at phylum plus Proteobacterial division and genus level classifications) and LEfSe analyses were performed on bacterial data sets rarefied to 59,000 sequences per sample and on fungal data sets rarefied to 10,000 sequences per sample, with each of these values being closest to the total filtered-sequence count of the smallest amplicon library (i.e. 59,540 sequences for the smallest bacterial 16S library and 10,716 sequences for the smallest fungal ITS2 library). The rarefaction was performed using the rarefy command in R package GUniFrac (Chen, 2018) on data matrices containing sequence counts from both sampling times.

### Hierarchical cluster analysis of (fungal and bacterial) phyla and abundant (top 20) genera composition between samples

Exploratory hierarchical cluster analysis was performed using average percent relative abundances of phyla (including Proteobacterial divisions for bacteria) and genera within sample type (N=6) using the Bray-Curtis dissimilarity index and average linkage method as performed in R version 3.1.2 (R Core Development Team, 2014) using commands in the Vegan (Oksanen et al., 2016) package and plotted with function Heatmap.2. For processed bacterial sequence data, we combined the sequence counts for genera *Afipia*, *Nitrobacter*, and *Bradyrhizobium* as an “NAB” genus category in the “top 20” gennera comparison. These genera were grouped together because they tend to have high sequence identity in their 16S rRNA-encoding genes, precluding robust discrimination of these genera with this marker sequence (Saito et al., 1998; Teske et al., 1994).

### LEfSe analyses

For both fungal and bacterial genus inventories, we used the LEfSe analysis tool (Segata et al., 2011) to make “within sampling time” comparisons of disturbance versus control plots within the same sampling depths (e.g. Comparison of Treatment A upper soil to Treatment C upper soil at 14 days).

For each LEfSe analysis, data matrices contained only sample units for two treatments compared against each other, and sequence counts per genus were converted to relative abundance of sequence totals per sample. Treatment category (A-D) was always used for the class comparison and plot replicates were used as subclasses for the Wilcoxon rank sum tests employed in the LEfSe algorithm. LEfSe analysis was run using the version downloadable at https://bitbucket.org/nsegata/LEfSe, with the default settings as follows: alpha value = 0.05 for the factorial Kruskal-Wallis test between classes, alpha value = 0.05 for the pairwise Wilcoxon test between subclasses, minimum logarithmic linear discriminant analysis score (LDA) score of 2 for inclusion as a differentiating genus, and 30 iterations for LDA bootstrapping. We report disturbance-indicating genera with p-values < 0.05 that are common to at least two treatments, because these genera likely represent microorganisms that tend to increase in abundance in the presence of detrital inputs from root disruption, hyphal disruption, or re-location of leaf litter. A complete inventory of disturbance indicators (as defined in the previous sentence), indicator LDA effect size scores, and box plots comparing relative abundance between sample types is provided in Supplementary Figures 3 (fungal) and 4 (bacterial).

## Results and Discussion

### Does fungal community composition distinguish soils from control plot versus disturbed plots?

Comparison of fungal phyla (Figure 1A) shows that 5-month post-disturbance, soils exhibit reduced relative abundance of ITS2 sequences assigned to phylum Basidiomycota and increased abundance for sequences assigned to phylum Zygomycota. Finer-scale comparison, focused on the twenty most abundant fungal genera reveals that 5-month post-disturbance soils cluster together because of reduced abundance Basidiomycete-associated EcM genera, alongside increased abundance for Zygomycete saprotrophs belonging to the genera *Mortierella* and *Umbelopsis* (Figure 1B). Concomitant saprotroph increases after disturbing soil EcM fungi were reported previously in ITS2 sequencing-based investigations (Kyaschenko et al., 2017; Lindahl et al., 2010; Stursová et al., 2014; Sukdeo et al., 2018). Newly disturbed samples (14-day sampling time) cluster with the control plots because of persistent, amplifiable ITS2 signatures from EcM. Specifically, the average relative abundances of *Amphinema*, *Cortinarius*, *Inocybe*, *Piloderma*, and *Sebacina* from disturbed samples are comparable to those of controls at 14 days (Figure 1B). This observation suggests that at 14 days post-disturbance, a large proportion of EcM detritus remains intact except at sites of severing, so EcM-derived ITS2 sequences are still amplifiable from soil DNA extracts. Increases in *Mortierella* abundance (via T-RFLP of amplified ITS regions) 14 days after root disruption in boreal forest humus layers have been reported previously (Lindahl et al., 2010), yet our results do not show strong differences in average *Mortierella* relative abundance between control and disturbed samples at the 14-day sampling time.

**Figure 1:**
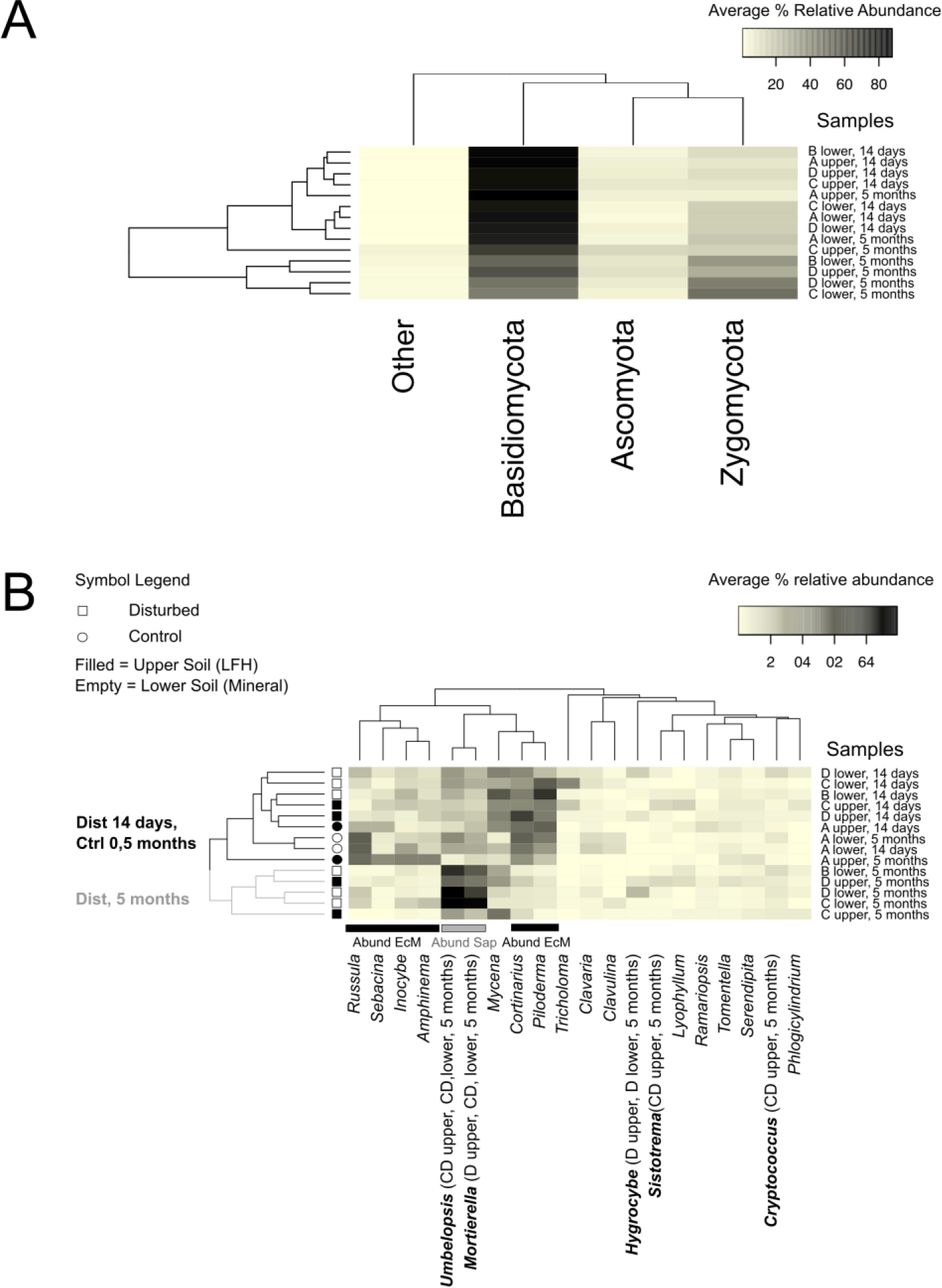
A) Hierarchical cluster analysis of fungal community composition at phylum level B) Hierarchical cluster analysis of 20 most abundant fungal genera in samples. Dominant EcM genera and disturbance-stimulated saprotrophic genera (Sap) that strongly influence sample clustering are indicated with labeled bars. Disturbance indicators (LEfSe-identified) are in bold with the associated disturbance regime and sampling time described in parentheses.

### Does bacterial community composition distinguish soils from control plot versus disturbed plots?

Comparison of bacterial 16S data at the phylum level for all samples and sampling times shows different community profiles for upper versus lower soils (Figure 2A). Lower soil samples contain a higher proportion of sequences assigned to phyla Acidobacteria, Gemmatimonadetes, and Verrucomicrobia compared to upper soils, while upper soils contain higher proportions of Actinobacterial sequences. In contrast to the conspicuous disturbance effects on fungal community composition, bacterial phyla and dominant genera profiles do not show separation of disturbed samples from controls, or separation by sampling times (Figure 2B). The minimal response of boreal forest humus-associated bacteria to root severing disturbances in contrast to strong response by fungal saprotrophs was reported by Lindahl, De Boer, and Finlay (Lindahl et al., 2010), and they suggested competitive exclusion by fast growing fungal saprotrophs as a reason for this disparity. Further, pyrotag sequencing of soils from logging treatments across multiple British Columbian forest ecozones indicated smaller responses of bacterial communities compared to fungal communities (Hartmann et al., 2012). Hartmann et al. suggest that forest soil bacteria may be more tolerant of deforestation-associated soil disturbances (including changes in moisture, temperature and aeration) and that their persistence in soil may rely less on symbiotic associations akin to EcM fungi interactions with dominant tree species within sites. Soil sampling in the Hartmann et al. study occurred 10-15 years after timber and forest floor removal events, in contrast to the shorter post-disturbance sampling times in our field study. Figure 2B shows multiple genera identified through LEfSe analyses, as disturbance indicators within both sampling times. This result suggests that we captured short-term (i.e. occurring over days to months) growth responses to detrital substrates, consistent with substrate-induced responses in other studies of litter decomposition (Bailey et al., 2017; Brabcová et al., 2016; Verastegui et al., 2014).

**Figure 2:**
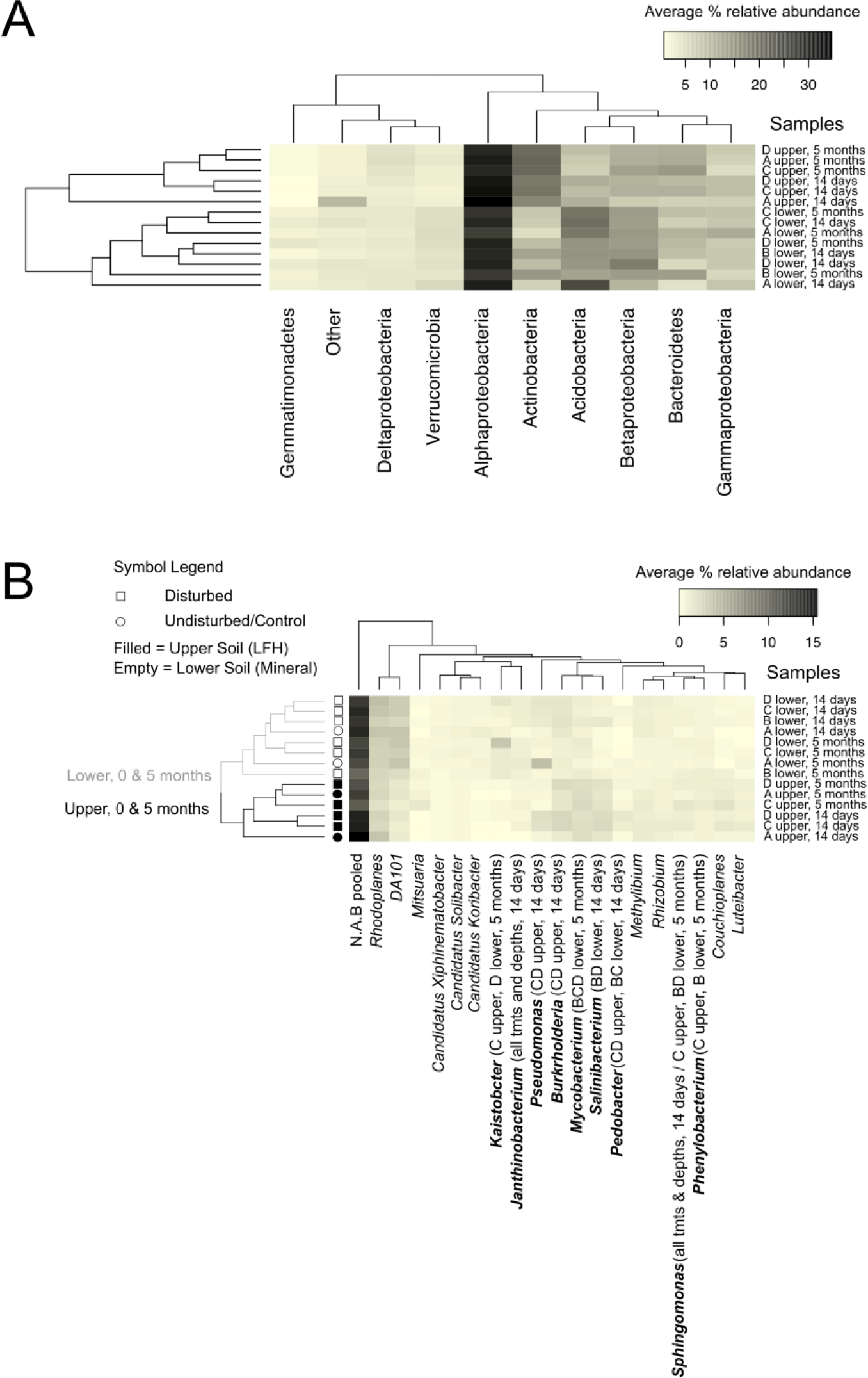
A) Hierarchical cluster analysis of bacterial community composition at phylum level with Proteobacterial divisions B) Hierarchical cluster analysis of 20 most abundant bacterial genera in samples. Disturbance indicators (LEfSe-identified) are in bold with the associated disturbance regime and sampling time described in parentheses.

### What is the strongest community composition feature that distinguishes physically disturbed soils from controls?

Hierarchical cluster analysis of abundant fungal genera alongside LEfSe comparisons of disturbed (Treatments B, C, and D) to control (Treatment A) plots show that soil mechanical disturbance strongly diminishes the abundance for EcM genera belonging to the class Agaricomycetes (Figure 3, Figure 1B). The highest observed LDA effect sizes (range: 4.74 – 5.25, considering both fungal and bacterial LDA scores) recovered from disturbed versus control plot comparisons by LEfSe, correspond to fungal class Agaricomycetes, fungal family Atheliaceae, and the EcM genus, *Russula* (Figure 3). Agaricomycetes indicates the control plots with LDA effect score sizes > 5 for all disturbance versus control comparisons at both sampling depths at 5 months. The Agaricomycete genera *Cortinarius*, *Russula*, and *Piloderma* exhibit reduced abundance across all disturbance regimes and both sampling depths at 5 months, and *Russula* exhibits disturbance-associated declines at 14-days post-disturbance. EcM Agaricomycetes genera like *Inocybe*, *Amphinema* and *Sebacina* exhibit diminished relative abundance in disturbed plots relative to controls at 5 months for upper soil samples, but not lower soil samples. These results suggest distinct distributions for different EcM taxa in our study site, such that *Cortinarius*, *Piloderma* and *Russula* are sensitive to soil disturbance across LFH to mineral horizon depths, whereas *Inocybe*, *Amphinema* and *Sebacina* preferentially colonize LFH materials. Because LEfSe retrieves class Agaricomycetes as a strong indicator for control plots in our study, the damage to EcM-derived biomass generates detrital inputs that become decomposition hot spots in disturbed plots.

**Figure 3:**
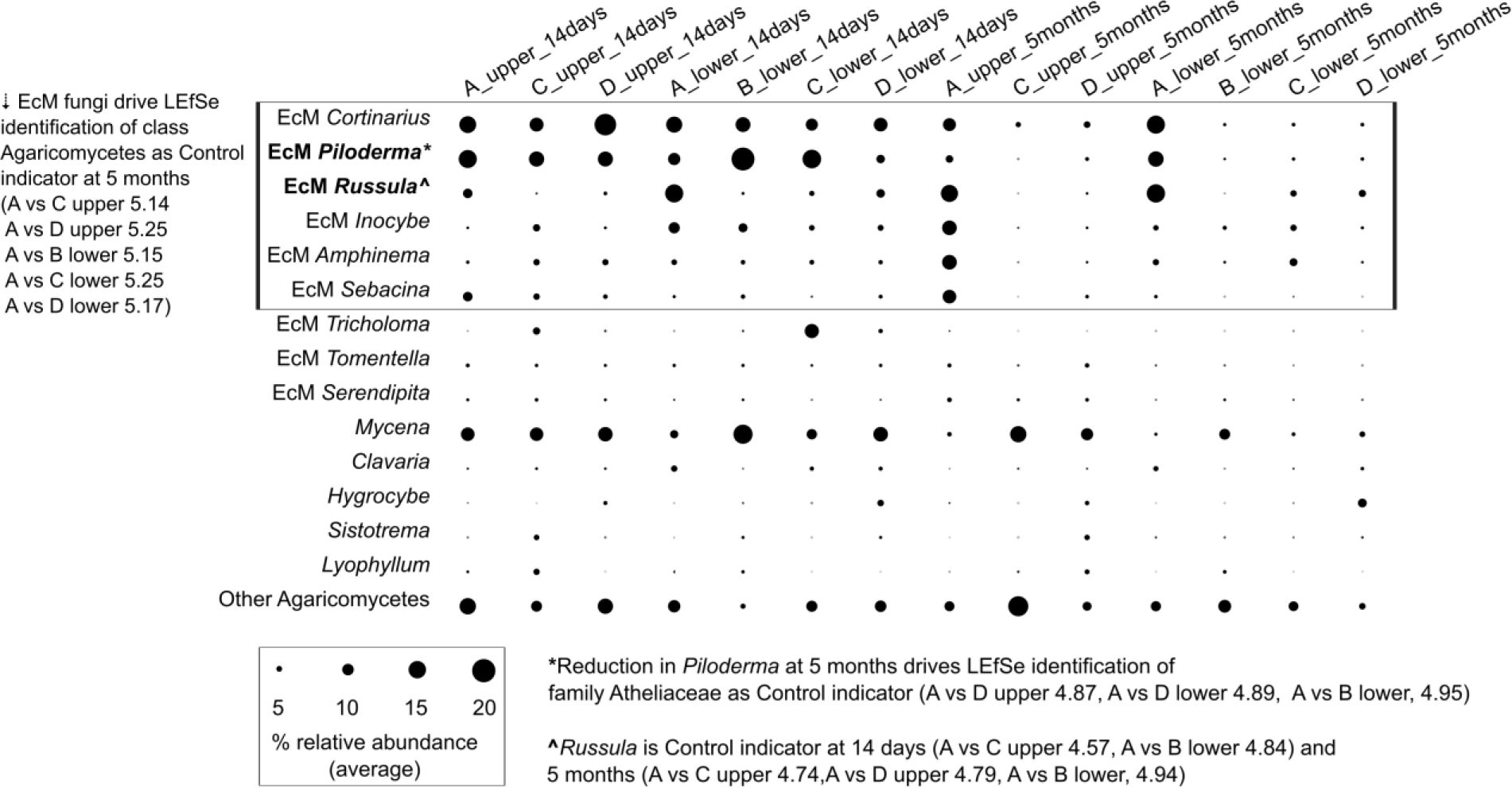
Bubble plot showing compositional shifts for the 14 most abundant fungal genera belonging to class Agaricomycetes in disturbed and control samples. Bubble radii are scaled to average percent relative abundance of each genera/category for rarefied sequence counts (10,000 reads/sample). Genera with known ectomycorrhizal guild associations are indicated with EcM.

### Do proxy fungal biomass measurements (ergosterol) suggest mycelial necromass turnover in disturbed soils?

We previously reported higher fungal biomass in upper soils from control plots compared to Treatments C and D, with statistically significant differences in distributions of ergosterol levels detected for A versus D comparison (Chi-square = 7.7, p = 0.021, with p_A-D_ = 0.024, Supplementary Table 1) (Sukdeo et al., 2018). This result mirrors observed reductions in EcM genera abundance in Treatment C and D upper soils sampled at 5 months post-disturbance (Figure 3).

For lower soil samples we previously reported significantly higher distributions of ergosterol values for Treatment B compared to Treatment A or D (Chi-square = 15.3, p = 0.002 with p_A-B_ = 0.003 and p_D-B_ = 0.005, Supplementary Table 1) (Sukdeo et al., 2018). Higher ergosterol content in Treatment B plots compared to lower soils from the other treatments may indicate the addition of LFH-associated fungi from horizon mixing, or saprotrophic fungal growth in response to multiple detrital types introduced by mixing. Lower soil samples contain less ergosterol on average compared to upper soils, yet declines in EcM genera are observed for disturbed soils from both sampling depths (Figure 3). These features of the ergosterol and ITS2 sequencing data establish that disturbed plots are a distinct substrate pool from control plots because of rapidly consumed necromass originating from EcM fungi damaged by the disturbance regimes. Increases in disturbance indicators *Mortierella* and *Umbelopsis* at 5 months are major disturbance-associated shifts in fungal community composition (Figures 1B, 5). This is a marked contrast to the responses of disturbance-indicating bacterial genera where abundance increases are detectable only 14-days post-disturbance, and do not increases in relative abundance enough to differentiate disturbed plots from controls. Therefore, mycelial and root necromass from mechanical disturbance stimulate fungal saprotrophs with different intensity and timing compared to bacterial saprotrophs.

### Do soil hydrolase pools increase with disturbance-associated root and mycelial necromass inputs?

The observed range of potential NAG, PHOS, and CBH activities in upper soil samples do not significantly differ between disturbed and control plots (Supplementary Figure 1) at either sampling time (14 days: Chi-square = 2.67 and p = 0.26 for NAG, Chi-square = 4.88 and p = 0.09 for PHOS, Chi-square = 1.20 and p = 0.55 for CBH, 5 months: Chi-square = 0.85 and p = 0.65 for NAG, Chi-square = 2.85 and p = 0.24 for PHOS, Chi-square = 1.91 and p = 0.39 for CBH). These results suggest saprotroph-derived hydrolases are not extensively secreted throughout disturbed soils containing fresh root and fungal necromass inputs. This result aligns with recent observations of higher levels of enzyme activity on buried *Tylopilus felleus* necromass compared to activities from surrounding forest soil or litter (Brabcová et al., 2016). Further, gel zymography studies indicate that severed roots (Spohn and Kuzyakov, 2014) and fungal mycelia (Guhr et al., 2015) exhibit maximal potential hydrolase activities when assayed directly at the hotspot location, so detecting net increases in activity at detrital hotspots by assaying bulk soil suspensions is not favorable. Stockpiling LFH (Treatment D) does not appear to impact the range or persistence of bulk soil-associated hydrolase activities when compared to immediately-replaced or undisturbed LFH (Supplementary Figure 1), suggesting that this reclamation strategy is minimally impactful on reservoirs of forest soil hydrolases.

For lower soil samples, ranges of observed NAG and PHOS activities but not CBH activity are significantly different between disturbance regimes (Supplementary Figure 2) 14 days post-disturbance (Chi-square = 10.29 and p = 0.02 for NAG, Chi-square = 13.79 and p = 0.003 for PHOS, Chi-square = 2.1 and p = 0.55 for CBH). Post hoc comparisons indicate significantly different soil potential NAG activity for Treatment B versus A (pA-B = 0.017), as well as for soil potential PHOS activities, comparing B versus A (pA-B = 0.01) and B versus D (pB-D = 0.005) comparisons. At 5 months post-disturbance, statistically significant differences for lower soil potential hydrolase activity (Supplementary Figure 2) are detected only for CBH levels, specifically between Treatments A and B (Chi-square = 9.25 and p = 0.03, pA-B = 0.02). Consistent with previous studies of forest soil enzyme activities (Baldrian et al., 2012), we notice higher levels of all measured potential hydrolase activities in upper versus lower soils across all treatments, mirroring the results of Phillips, Ward, and Jones (Phillips et al., 2014) that soils with abundant fungal biomass from saprotrophic or EcM inhabitants exhibit comparable hydrolytic enzyme activities. All disturbed lower soils exhibit higher and more variable potential NAG activity at 14 days post-disturbance compared to control soils (Supplementary Figure 2). We do not see a similar trend for disturbed versus control upper soils (Supplementary Figure 1), which have higher fungal biomass and higher, more variable NAG activity levels in general. Therefore lower soils might exhibit higher chitinase activities post-disturbance since mycelial necromass hotspots are more evenly distributed by mixing compared to discrete areas of detrital turnover in control plots. The highest observed values for potential NAG and PHOS at 14 days and for CBH at 5 months in Treatment B (compared to other lower soil samples) is expected given the incorporation of LFH material into underlying mineral soil.

### What ecological functions are associated with disturbance-indicating fungal genera?

*Capronia* was retrieved as a disturbance indicator (Figure 4) common to three or more sample types (i.e. a sample group defined by disturbance regime and sampling depth) at 14 days. The fungal genus *Capronia* has been previously reported as an early responder to EcM/root-severing in forest humus (Lindahl et al., 2010) and this is consistent with retrieving this genus as an indicator for multiple disturbance regimes. Recent DNA stable-isotope-probing (DNA-SIP) investigations of North American forest soils suggest that members of the genus *Capronia* are hemicellulose degraders (Leung et al., 2016), which would be relevant to invasion and/or colonization of severed plant roots.

At 5 months post-disturbance, 10 disturbance-indicating genera common to three or more sample types were retrieved through LEfSe analysis (Figure 4). Our analysis did not recover any fungal genera that indicated disturbance common to all sample types but *Rhinocladiella*, *Umbelopsis*, and *Volutella* were retrieved as disturbance indicators common to four sample types (Figure 4). Consistent with hierarchical clustering analysis results (Figure 1B), *Mortierella* and *Umbelopsis* exhibit the highest observed LDA scores for control versus disturbance regime comparisons, all of which were greater than 4.5 (Figure 4). *Hygrocybe* was retrieved as a disturbance indicator for Treatment D, upper and lower soil samples with an LDA score > 4. Fungal disturbance indicators detected at 14 days and 5 months were *Capronia* and *Trichoderma* (Figure 4). Upper soil samples of disturbed plots (i.e. Treatments C and D, upper soils) should contain the largest inputs of mycelial necromass due to losses of EcM taxa (Figure 3), and the fungal disturbance indicators common to both of these treatments include *Cryptococcus*, *Mastigobasidium*, *Mucor*, *Sistotrema*, *Trichoderma*, and *Umbelopsis* (Figure 4).

Fahad (2017) used RNA-SIP to identify fungal inhabitants of boreal forest podzols that consume ^13^C-labelled *Piloderma fallax* mycelia and observed enrichment of saprotrophic genera including *Mortierella*, *Mucor*, *Penicillium*, *Trichoderma*, and *Umbelopsis*. The results of this RNA-SIP investigation support our findings that these genera, particularly *Mortierella* and *Umbelopsis*, are stimulated by fungal detritus and remain abundant for some time after necromass pools are depleted from soils (Fahad, 2017). Brabcovà *et al.* (2016) investigated fungal decomposers in litter and soils from a temperate oak forest litter and soils using *Tylopilus felleus* sporocarp tissue buried in either horizon. Their amplicon sequencing results showed enrichment of fungi belonging to the genera *Penicillium*, *Aspergillus*, *Crocicreas*, *Tomentella*, *Epicoccus*, *Mucor*, and *Candida* on *T. felleus* tissue buried in litter, while *Mortierella*, *Aspergillus*, *Cladosporium*, *Mucor*, and *Kappamyces* were enriched on mycelial necromass buried in soil. We detect increased abundance of *Mucor* and *Mortierella* sequences in disturbed upper and lower soil samples at 5 months (Figure 4) suggesting saprotrophic colonization of mycelial detritus across the 20 cm depth of our disturbance regimes.

In our study, disturbed plots contained pools of labile necromass in the form of severed roots and LFH litter that were incorporated into lower soil horizons from incidental mixing when disturbance regimes were applied. Disturbance indicators (Figure 5) previously associated with decomposing leaf litter include *Cryptococcus* (Štursová et al., 2012; Voříšková and Baldrian, 2013), *Mortierella* (Štursová et al., 2012; Voříšková and Baldrian, 2013), and *Sistotrema* (Štursová et al., 2012). We retrieved other disturbance indicators with potential plant necromass associations such as needle litter colonization (*Volutella*) (Osono and Takeda, 2007), root pathogenicity (*Entoloma*) (Gryndler et al., 2010; Hietala and Sen, 1996) and wood decay (*Rhinocladiella*) (Lumley et al., 2001).

**Figure 4:**
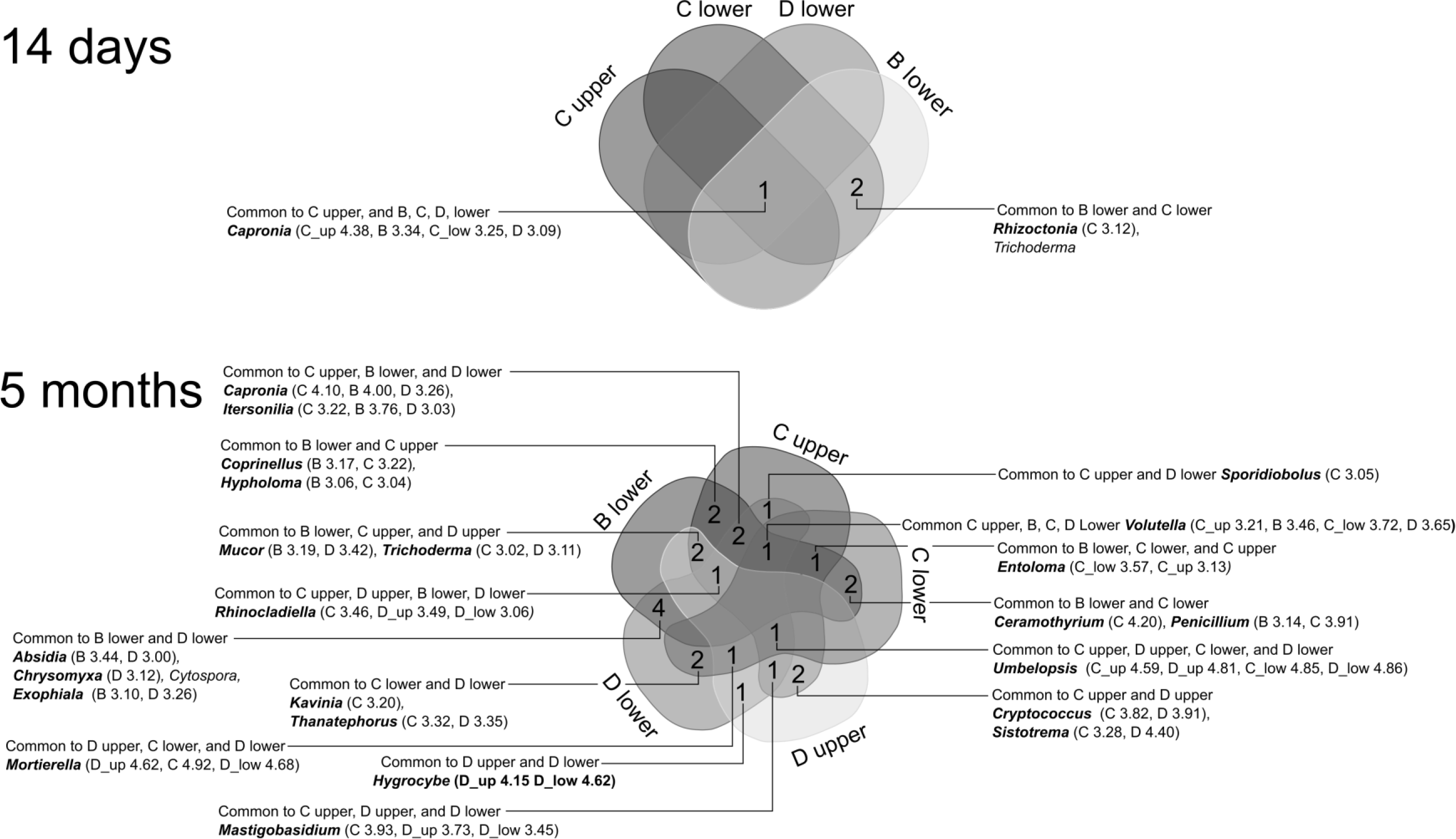
Venn diagram shows disturbance-indicating fungal genera for 14-day and 5-month sampling times. Genera names in bold have LDA scores > 3 for at least one treatment versus control comparison and scores with values > 3 are listed in parentheses.

### Which bacterial genera distinguish soils from disturbed plots versus controls?

LEfSe comparison of disturbed versus control plots at 14 days retrieved 24 bacterial disturbance indicator genera (Figure 5) common to multiple sample types (in contrast to only 3 fungal disturbance indicators). Two bacterial disturbance indicators, *Janthinobacterium* and *Sphingomonas*, are common to all disturbance treatments and sampling depths. LDA effect sizes for *Janthinobacterium* are slightly higher in disturbed lower soil samples but higher in upper soil samples for *Sphingomonas* (Figure 5). The highest observed LDA effect sizes (score > 3.7) for bacterial genera at 14 days were retrieved for *Burkholderia* and *Pseudomonas*, both of which are disturbance indicators for Treatments C and D upper soils, and *Pedobacter*, which indicates disturbance in Treatment D upper soil (Figure 5).

LEfSe comparisons for bacterial community profiles at the 5-month sampling time retrieved disturbance indicators for all comparisons except Treatment D versus A, upper soil. Seven disturbance indicators retrieved at 14 days (*Aeromicrobium*, *Cellvibrio*, *Hymenobacter*, *Pseudonocardia*, *Rhodoferax*, *Sphingomonas* and *Virgisporangium*) persist as disturbance indicators for multiple sample types at 5 months (Figure 5), and all of these genera are disturbance indicators for Treatment B in the later sampling time.

**Figure 5:**
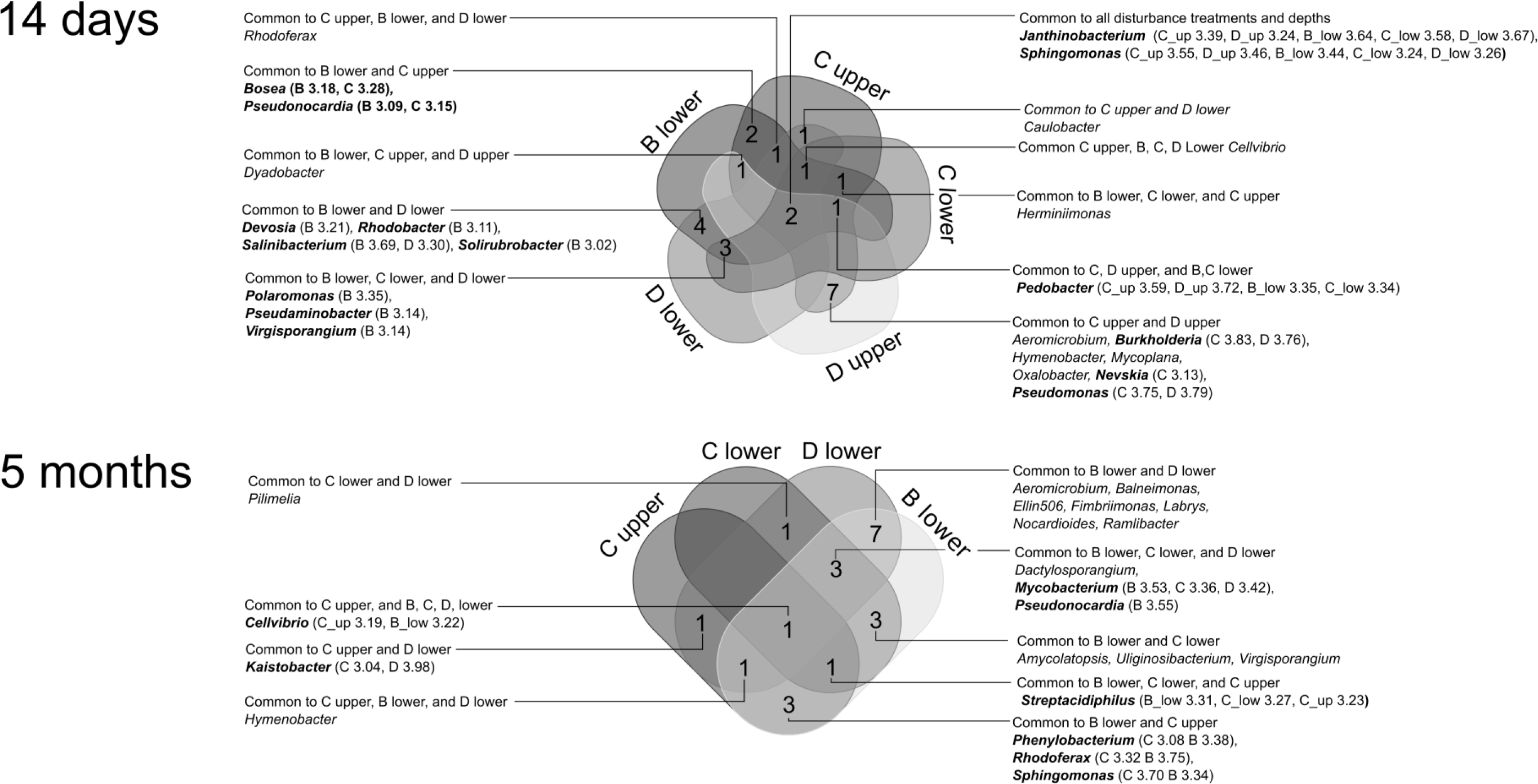
Venn diagram shows disturbance-indicating bacterial genera for 14-day and 5-month sampling times. Genera names in bold have LDA scores > 3 for at least one treatment versus control comparison and scores with values > 3 are listed in parentheses.

### Are bacterial disturbance indicators potential responders to fungal necromass?

Several disturbance indicators (*Burkholderia*, *Janthinobacterium*, *Pedobacter*, and *Pseudomonas*) from our study are genera that Brabcová *et al.* (2016) identified as abundant colonizers of *T. felleus* necromass (Supplementary Table 2) buried in oak forest litter or soil. *Sphingomonas*, a general disturbance indicator in our field study at 14 days, increased in abundance over time on *T. felleus* necromass buried in soil, and *Phenylobacterium* (disturbance indicator for C upper soil and B lower soil at 5 months) increased in abundance on necromass buried in litter at later time points, suggesting that members of these genera are also responsive to fungal necromass for growth substrates (Brabcová et al., 2016). In Fahad’s RNA-SIP study of bacterial and fungal *P. fallax* assimilation, the isotopically enriched community profile was characterized 28 days after introduction of labeled mycelium to soil microcosms (Fahad, 2017). Although our study profiles microbial community shifts at 14 days and 5 months after the creation of necromass inputs, there is overlap between disturbance-indicating genera we identify, and those identified by Fahad as consumers of *P. fallax* detrital materials. These overlapping genera include *Burkholderia*, *Herminiimonas*, *Pedobacter*, and *Streptacidiphilus* (Supplementary Table 2). Our results demonstrate that we can identify fungal necromass-responsive bacteria by 16S amplicon sequencing of bulk soils if they are sampled 14 days post-disturbance.

Fungal tissue contains chitin and chitosan as major structural components making it a substrate distinct from plant litter. Because of the abundant EcM biomass in forests, turnover of EcM derived substrates can be expected to stimulate bacteria that respond to the presence of chitin. Debode *et al.* observed increases in lettuce rhizosphere soil abundance of *Cellvibrio*, *Pedobacter*, and *Sphingomonas* with 2% chitin amendments to potting soil (Supplementary Table 2), compared to un-amended controls (Debode et al., 2016). These genera had some of the highest fold increases in abundance with chitin amendment and this mirrors our results where *Cellvibrio*, *Pedobacter*, and *Sphingomonas* exhibit some of the highest bacterial indicator LDA scores (i.e. >3.5 for *Pedobacter* and *Sphingomonas*, Supplementary Figure 4). Other genera stimulated by chitin (Debode et al., 2016) that are more abundant in our disturbance treatments include *Nocardioides* and *Dyadobacter* (Supplementary Table 2).

Bai (2015) observed strong increases in amplifiable *Janthinobacterium*-associated chitinase A sequence abundance for forest and grassland soils incubated with various chitin substrates (derived from shrimp, *Mucor hiemalis*, *Aspergillus niger*, and mealworm). Bacterial 16S pyrosequencing in Bai’s research recovered abundant sequences belonging to family Pseudomonadaceae from chitin samples incubated in forest soil (Bai, 2015). Our observations of disturbance-associated increases in abundance for *Janthinobacterium* and genera from Pseudomonadaceae (specifically *Pseudomonas* and *Cellvibrio)* suggests similarities between bacterial responses to exogenous chitin inputs in soils and chitin inputs from *in situ* fungal necromass pools.

### Are bacterial disturbance indicators associated with other detrital substrates?

The disturbance regimes in our field study created extensive pools of root necromass and relocated litter material at different depths in a 20 cm soil profile compared to control plots. Such detrital inputs provide bioavailable nutrient pools that stimulate bacterial heterotrophs, but they also represent substrate contexts that recruit bacteria to rhizosphere or mycorrhizosphere locations where they perform ecosystem services relevant to such soil compartments. Verastegui et al. (2014) suggest, in their investigation of soil bacterial carbohydrate utilization by DNA-SIP, that labeled substrate concentrations in incubations are representative of soil compartments associated with rhizosphere or plant material decomposition processes, since they were higher than typical bulk soil concentrations. Therefore we can consider how short-term bacterial responses to litter-derived substrates in SIP studies (including RNA, DNA, and PLFA SIP), plant-derived litter, and mycorrhizospheric interfaces may promote localized abundances of genera that we identify as disturbance indicators.

*Cellvibrio*, a broadly detected disturbance indicator in our study, has been associated with cellulolytic and hemicellulolytic activity (Supplementary Table 2) (Wilhelm, 2016), and increased abundance in soils with labelled root and leaf litter amendments (Kramer et al., 2016). In addition to reported chitinolytic activity of members of genus *Janthinobacterium* (Kielak et al., 2013), hemicellulose enrichment of this genus in SIP investigations (Leung et al., 2016; Wilhelm, 2016) suggests multiple litter decomposition functions for this disturbance indicator. Bacteria belonging to the genus *Burkholderia*, are described as having beneficial associations with mycorrhizal fungi and broadly associated with mycrorrhizoplane/mycorrhizosphere soil compartments (Marupakula, 2016). *Burkholderia* has been observed a strong responder to hemicellulose additions in forest soils (Leung et al., 2016), and previously identified as an enriched genus in DNA-SIP experiments using arabinose, cellobiose, cellulose, glucose, and xylose amendments in soil (Štursová et al., 2012; Verastegui et al., 2014). The disturbance indicators *Herminiimonas*, *Pedobacter*, *Sphingomonas*, and *Streptacidiphilus* (Supplementary Table 2) have been identified as genera stimulated by cellulose inputs (Schellenberger et al., 2010; Štursová et al., 2012; Wilhelm, 2016).

Several disturbance indicator genera (*Aeromicrobium*, *Devosia*, *Labrys*, *Ramlibacter*, *Phenylobacterium*, and *Salinibacterium*) were retrieved from lower soil samples, particularly from Treatment B and Treatment D. Verastegui et al. (2014) observed a broad response of these genera to multiple plant-associated substrates by DNA-SIP (Supplementary Table 2). These results suggest that poorly reconstructed soil profiles (i.e. Treatment B) and exposed mineral soils (i.e. Treatment D, lower soil) contain bacterial inhabitants capable of detritus turnover, but LDA effect sizes for genera are generally lower than those observed for *Burkholderia*, *Cellvibrio*, *Janthinobacterium*, *Pseudomonas*, and *Sphingomonas*, or upper soil disturbance indicators like *Burkholderia* and *Pseudomonas*.

## Conclusions

Strong reduction of EcM fungal genera was detected in all disturbed plots at 5-months post-disturbance compared to controls (Figure 6), which adds to prior evidence for the diagnostic value of ITS2 sequencing for monitoring mycorrhizal decline and/or recovery in forest soils with a histories of natural or anthropogenic disruption. Although strong increases in saprotrophic fungal genera such as *Mortierella* and *Umbelopsis* were observed across multiple disturbance regimes, soil potential hydrolase activities did not suggest a uniform, intensive increase in bulk soil enzyme pools (Figure 6) associated with organic matter degradation. Reductions in EcM genera coincided with reduced fungal biomass (i.e. ergosterol) in all disturbed upper soils (Figure 6), providing an opportunity to compare bacterial and fungal responses distributed pools of mycelial necromass within five months. Soil collected from disturbed plots after 14-days exhibited enrichment of bacterial genera associated with mycelial necromass (i.e. *Burkholderia*, *Cellvibrio*, *Janthinobacterium*, *Pedobacter*, *Pseudomonas,* and *Sphingomonas*) and with enrichment following cellulose inputs (i.e. *Burkholderia*, *Cellvibrio*, *Pedobacter*, and *Sphingomonas*).

These results demonstrate that bacterial 16S profiling of bulk soils in our study captures rapid, short-lived enrichment of bacterial saprotrophs following mycelial and root necromass inputs from disturbance (Figure 6). Therefore we suggest that adding diagnostic value to bacterial 16S surveys in soils may rely on sampling schemes that follow transient incidental or deliberate stimulation of dynamic processes, like organic matter degradation, to evaluate whether process presence, rates, and corresponding community composition changes differ according to the soil disturbance(s) under investigation. Leung et al.’s (Leung et al., 2016) investigation of hemicellulose-stimulated (i.e. DNA-SIP) microbial (fungal and bacterial) communities in post-harvest forest soils exemplify utility of this approach. Although hemicellulose-induced respiration rates did not significantly differ between post-harvest soils with three different levels of organic matter removal, the population structures of hemicellulose-stimulated bacteria and fungi taxa were distinct between control soils when compared to post-harvest soils where forest floor material was also removed. DNA-SIP investigations of LTSP soil samples using cellulose have also revealed differences in cellulolytic fungi and bacterial abundances between control and harvest plus organic matter removal treatments (Wilhelm et al., 2017). Therefore artificial (i.e. SIP) or incidental (i.e detritusphere creation from disturbance) substrate stimulation in experimental designs is beneficial for generating response-based categories of bacterial and fungal saprotrophs from amplicon sequencing data. These saprotrophic guilds could be useful for relating forest soil disturbance histories to resulting shifts substrate-induced population structures attributable to modified densities, turnover rates, and distributions of detritusphere, rhizosphere and ectomycorrhizal compartments.

**Figure 6:**
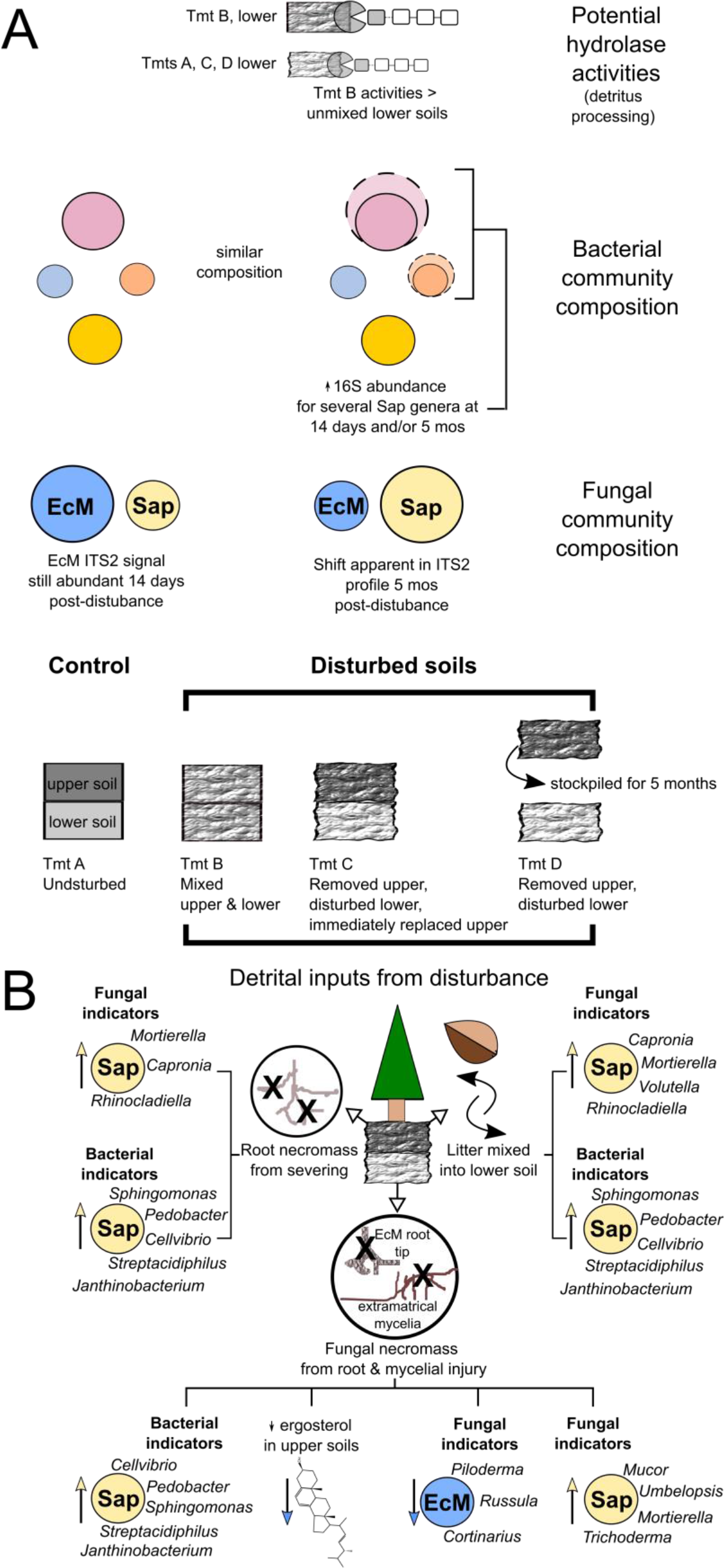
Summary of field study disturbance regimes (between thick black brackets), A) Observed disturbance effects on potential hydrolase activities, bacterial community composition, and fungal community composition, and B) Summary of disturbance-associated detrital inputs and saprotrophic (Sap) disturbance indicator genera grouped by detritusphere associations described in the research literature. Fungal biomass loss (i.e. ergosterol) and ectomycorrhizal (EcM) genera declines are also included as indirect evidence of a labile fungal necromass pool created through soil disturbance. Indicator genera shown are those with LDA scores > 3 that were retrieved in three or more disturbance regime versus control comparisons by LEfSe analysis. Circles representing relative abundances of Sap fungi, EcM fungi, and bacterial genera are not to scale.

## Acknowledgements

We thank Jessica R. Lowry and Jennifer Salokannel for preparing ready-to-sequence amplicon libraries. We are grateful that Clive Dawson and other laboratory staff at the Analytical Chemistry Laboratory, B.C. Ministry of Environment, Knowledge Management Branch, performed chemical analysis of soil samples for this study. We thank the Génome Québec Innovation Centre for high throughput sequencing services and bioinformatics analysis of sequencing data. We specifically acknowledge the Génome Québec Innovation Centre personnel as follows: Dr. Alfredo Staffa for technical guidance relating to amplicon preparation, as well as Dr. Pascale Marquis and Dr. Mathieu Bourgey for completing the bioinformatics analysis of amplicon sequencing data. We are very grateful to Dr. Michael Gillingham and Dr. Jeanne Robert for financial, administrative, and logistic support during the duration of this research project.

